# An integrated gene catalog of the zebrafish gut microbiome reveals significant homology with mammalian microbiomes

**DOI:** 10.1101/2020.06.15.153924

**Authors:** Christopher A. Gaulke, Laura M. Beaver, Courtney R. Armour, Ian R. Humphreys, Carrie L. Barton, Robyn L. Tanguay, Emily Ho, Thomas J. Sharpton

**Affiliations:** Department of Microbiology, Oregon State University, Corvallis, OR 97331, United States; School of Biological and Population Health Sciences, Oregon State University, Corvallis, OR 97331, United States; Department of Environmental and Molecular Toxicology, Oregon State University, Corvallis, OR 97331, United States; Linus Pauling Institute, Oregon State University, Corvallis, OR 97331, United States; Department of Statistics, Oregon State University, Corvallis, OR 97331, United States

**Keywords:** microbiome, metagenome, intestine, vertebrate, zebrafish, diet

## Abstract

Gut microbiome research increasingly utilizes zebrafish (*Danio rerio*) given their amenability to high-throughput experimental designs. However, the utility of zebrafish for discerning translationally relevant host-microbiome interactions is constrained by a paucity of knowledge about the biological functions that zebrafish gut microbiota can execute, how these functions associate with zebrafish physiology, and the degree of homology between the genes encoded by the zebrafish and human gut microbiomes. To address this knowledge gap, we generated a foundational catalog of zebrafish gut microbiome genomic diversity consisting of 1,569,102 non-redundant genes from twenty-nine individual fish. We identified hundreds of novel microbial genes as well as dozens of biosynthetic gene clusters of potential clinical interest. The genomic diversity of the zebrafish gut microbiome varied significantly across diets and this variance associated with altered expression of intestinal genes involved in inflammation and immune activation. Zebrafish, mouse, and human fecal microbiomes shared > 50% of their total genomic diversity and the vast majority of gene family abundance for each individual metagenome (∼99%) was accounted for by genes that comprised this shared fraction. These results indicate that the zebrafish gut houses a functionally diverse microbial community that manifests extensive homology to that of humans and mice despite substantial disparities in taxonomic composition. We anticipate that the gene catalog developed here will enable future mechanistic study of host-microbiome interactions using the zebrafish model.

**Importance:** Zebrafish have emerged as an important model system for defining host-microbiome interactions. However, the utility of this model is blunted by limited insight into the functions that are carried by zebrafish gut microbiota, their relationship with zebrafish physiology, and their consistency with the functions carried by human gut microbiota. To address these limitations, we constructed the first genomic database of zebrafish gut microbiome diversity. We use this novel resource to demonstrate that the genomic diversity of the zebrafish gut microbiome varies with diet and this variance links with altered intestinal gene expression. We also identify substantial homology between zebrafish, human, and mouse metagenomic diversity, indicating that these microbiomes may operate similarly.

## Introduction

Overwhelming evidence has identified the human gut microbiome as a crucial element of health^1–3^. Yet, our knowledge of the specific mechanisms through which the microbiome elicits effects on health remain obscure. To move beyond describing the association between microbiome composition and disease and towards elucidating these mechanisms, researchers need robust tools and experimental systems that can accurately measure and model these relationships. Animal model systems, especially the mouse model, have proven critical in this regard and clarified specific mechanisms underpinning of host-microbiome interactions^4^. But because of the overwhelming genetic diversity of the gut metagenome^5^ and the staggering set of factors that may influence microbiome operation, researchers need access to animal models that can accommodate high-throughput screening in the effort to identify stimuli that shape the microbiome’s associations with health.

The zebrafish is an emerging high-throughput animal model system of host-microbe interactions. The high fecundity, short generation time, transparent life stages, and simple gastrointestinal tract makes zebrafish an ideal model for studying the fundamental molecular interactions between hosts and their microbiomes. Foundational research using this model system identified conserved immune responses to gut microbes between zebrafish and mammals^6^. Other reports indicated that the microbiomes of mammals could colonize the zebrafish gut and vice versa, hinting that the microbiomes of these animals may perform similar ecological functions in the gut^7^. Conversely, compositional differences between mammalian and zebrafish microbiomes are significant^8^ and argue that there may exist substantial functional differences between mammalian and zebrafish microbiomes. However, little is known about the genetic diversity of the zebrafish gut microbiome, which hampers our ability to adequately assess the functional similarity between the zebrafish and human gut microbiomes.

Shotgun metagenomic sequencing is a recently emerged method for characterizing the functional capacity of the gut microbiome^9^. In particular, metagenomes facilitate the generation of integrated gene catalogs (IGCs)^10^, which document the specific genes, protein families, and molecular functions encoded in the gut microbiome. These vital data resources have advanced understanding of how the functions employed by gut microbes link to human health as well as the consistency in functional capacity between human and non-human gut microbiomes^11,12^. However, metagenomes have, to date, not been capitalized upon as a zebrafish microbiome research method.

Here, we cataloged the genomic diversity of the zebrafish gut microbiome and provide the first detailed insights into the operation of zebrafish gut microbial communities. We demonstrate that the zebrafish gut microbiome functional potential varies between fish fed defined laboratory diets and that this variance associates with altered zebrafish intestinal gene expression. Finally, we used this gene catalog to quantify similarities between human, mouse, and zebrafish metagenomic diversity and identified substantial homology across these vertebrates.

## Methods

### Animal Subjects, Sample Collection, and Sequencing

This study was conducted in accordance with the recommendations in the Guide for the Care and Use of Laboratory Animals of the National Institutes of Health and with protocols approved by the Oregon State University Institutional Animal Care and Use Committee (IACUC). Forty-five healthy adult, male, zebrafish (5D) were fed one of three diets (Supplemental table 1): 1) a standard lab diet (LDC) (Gemma Micro. Skretting, Tooele, France; N=15), 2) a defined diet (DDC) with moderate amounts of zinc (34mg/kg; N=15), and 3) a defined diet (CZMD) with low levels of zinc (14mg/kg; N=15). Our previous work demonstrated that the DDC diet contains physiologically adequate levels of zinc, while the levels of zinc in the CZMD diet induce marginal zinc deficiency in zebrafish^13^. Purified defined diets were prepared as previously described^13^. Zebrafish were group housed in three tanks per treatment group (N = 5 / tank) at a density of ∼6 fish / L. After nine months on the diet, the animals were individually isolated and housed overnight in 0.5L clean spawning tanks. Fecal materials were then collected from each individual fish’s tank using a sterile transfer pipette. These samples were then rapidly frozen on dry ice and stored at -20°C until processing. DNA was isolated from fish fecal samples using the MOBIO PowerSoil kit as previously described^14^. Shotgun metagenomic libraries were then constructed from a subset of these animals (LDC N = 15, DDC N = 7, CZMD N = 7) using the Nextera XT kit (Illumina, San Diego, CA USA) and sequenced on and Illumina HiSeq 3000 using a 150bp PE sequencing kit (Illumina).

### Metagenome Assembly and Functional Annotation

Each metagenomic library was demultiplexed and quality filtered with shotcleaner (https://github.com/sharpton/shotcleaner) using the zebrafish genome (GRCz10). The cleaned sequence files were then assembled simultaneously using MEGAHIT^15^ with a minimum kmer size of 27 and a kmer increment size of 10. The resulting contigs were used to predict genes using prodigal in metagenome mode. CD-HIT^16^ was used to collapse redundant genes (95% nucleotide identity, 90% coverage of the shorter sequence) into a singular representative sequence. An all vs. all local protein alignment was the conducted to identify duplicates using DIAMOND (v0.9.4)^17^. Alignments with > 90% protein coverage (shorter sequence) and 95% amino acid identity were considered duplicates and the shorter of the two proteins was subsequently removed. This non-redundant set of proteins was then aligned to the KEGG (v 73.1)^18^, EGGNOG (v4.5)^19^, Pfam^20^, and vFam (v2014)^21^ protein family databases using DIAMOND. Each predicted protein with an alignment passing a permissive filter threshold (e-value < 0.001) was annotated with its best hit. Gene family diversity was assessed using the R package vegan functions vegan::diversity and vegan::rarefy in R.

### Protein Family Clustering

Pairwise alignments between predicted zebrafish, human^10^, and mouse^12^ proteins were generated using DIAMOND (v0.9.4). Expect values were scaled to compensate for the total combined length of the three protein databases. Alignments were filtered to remove hits with coverage of < 80% of the longer sequence or e-value > 1×10^−10^. Clusters of homologous proteins were then generated using the Markov Cluster (MCL) algorithm^22^ with an inflation value of 2. This value was chosen as it has been demonstrated to be appropriate for clustering of microbial protein families^23^. A cluster was considered present in a species’ metagenome if that cluster contained at least one protein derived from the IGC of that species. Global alignments of clustered proteins were generated for each cluster that contained at least one representative from each species using MUSCLE^24^. FastTree^25^was used to construct phylogenetic trees from each of these alignments and phylogenetic signal in MCL protein families was assessed using Pagel’s lambda^26^.

### Homology Analysis

Inter-species homologous proteins sequences were identified from pairwise alignments of full-length zebrafish, human, and mice predicted proteins generated using DIAMOND (max e-value 1×10^−3^). Intra-species protein homologs were identified by performing reciprocal pairwise alignments of the set of full-length proteins for each species. Alignments were subsequently filtered to remove hits with < 80% coverage of the longer sequence.

### Genome Binning and Biosynthetic Gene Cluster Discovery

Metagenomic bins were generated using MaxBin 2.0 (v 2.2.1) with a minimum contig length of 1,000bp and a probability threshold of 0.9^27^. Completeness and contamination of each genome bin was calculated by assessing the percentage of the 107 universal single copy number genes present in each bin (completeness) and the percentage of all universal single copy number genes that contained multiple hits to a single copy number sequence in that bin (contamination)^28^. Coverage for each genome bin was computed using MaxBin 2.0 by aligning all sequence reads to the binned genomes using Bowtie 2^29^. Genome bin taxonomy was assessed using Kraken^30^and a bin was assigned a phylum label if > 50% of all contigs in that bin were associated with that taxonomic label.

Biosynthetic gene clusters (BGCs) were predicted for each genome bin using antiSMASH (v 3.0.5)^31^ with default parameters. Predicted BGC proteins were aligned to a protein database that contained all bacterial and archaeal proteins in RefSeq (v92)^32^ using DIAMOND. A single best-hit alignment was reported for each protein. Proteins for which no alignment met the e-value threshold (< 0.001) were classified as “no-hit”.

### RNA-Seq and Differential Gut Gene Expression Analysis

Whole intestinal tracts were collected at necropsy for each fish and stored at -80°C until processing. The intestinal tissue was homogenized (N = 7 fish / diet) using a pellet pestle in nuclease free microcentrifuge tubes and RNA was subsequently extracted from the homogenate using the RNeasy kit (Qiagen, Hilden Germany) according to the manufacturer’s instructions. Poly-A tailed RNA was enriched (PrepX; Takara Bio, Kasatsu Japan), strand specific RNA libraries (PrepX; Takara Bio, Kasatsu Japan) were constructed using an Apollo 324 robot (Takara Bio, Kasatsu Japan), and libraries were sequenced on an Illumina HiSeq. A total of 1.28×10^9^ 150bp paired end reads (mean 3.05×10^7^ PE reads /fish) were adaptor trimmed and quality filtered using ea-utils^33^. Quality controlled reads were then aligned to the zebrafish genome (GRCz11) using Tophat (v 2.1.1; options: max-insertion-length 5, max-deletion-length 5, read-gap-length 5, read-mismatches 5, read-edit-dist 10, mate-std-dev 50, min-anchor 5, splice-mismatches 1, b2-very-sensitive, b2-N 1)^34^. Transcript counts were calculated using HTSeq (v0.9.1, method = union)^35^. Global differences in gene expression between diets was calculated using a PERMANOVA (Euclidean distance). Differential gene expression between pairs of diets was calculated using DeSeq2 (Wald test)^36^ and false discovery rate was controlled using q-value^37^.

### Differential Microbial Gene Abundance Analysis

Sequencing reads were aligned to the IGC gene models using Bowtie 2. Differences in gene richness and Shannon entropy were quantified using a Kruskal-Wallis *H* test. An Adonis test (R; vegan) was used to quantify the variation in beta-diversity (Bray Curtis dissimilarity) attributable to differences in diet. Dietary differences in KEGG protein family, module, and pathway abundance was assessed using negative binomial generalized linear models (R; MASS) with a square root link function. Briefly, a single model that incorporated diet type as a parameter was constructed for each protein family, module, or pathway. Analysis of variance (ANOVA) was used to determine if the model that incorporated the diet parameter explained significantly more variation than a reduce model, which only included the intercept parameter. False discovery rate was controlled using q-value (R; qvalue).

### Correlation Network Analysis

To examine potential associations between dietary variance in microbiome function and zebrafish gut gene expression, we calculated Spearman’s rank order correlations between abundance of host genes and microbial pathway abundance that significantly varied across diet. Since host gene expression and microbial gene abundance data were collected in two distinct samplings at necropsy, mean tank abundances of expression and abundance data were used. As above, false discovery rate was controlled using q-value. Significant correlations (q < 0.2) were used to construct bipartite host-microbiome correlation networks. Degree and hub scores were calculated using igraph. Community analysis was conducted using the fast-greedy algorithm in igraph.

### Data Availability

All data used in these analyses have been deposited in the Sequence Read Archive under the project numbers PRJNA598878 and PRJNA598916.

## Results

Integrated gene catalogs have been widely used to elucidate links between microbiome operation and host physiology^11,38^, expand our knowledge of microbial functional and taxonomic diversity^10,12^, and identify novel drug leads^39^. We generated the first assembly of the zebrafish gut metagenome, which was derived from 29 zebrafish fecal metagenomes and consists of 1.06 x 10^9^ bp and 998,094 contigs. This assembly was similar in quality (N50 = 1,838bp, mean contig length = 1,064bp, median reads mapped to assembly = 97%) to the high-quality assemblies used to construct other vertebrate integrated gene catalogs including human^10^ and mouse^12^ IGCs. The assembled contigs contained 1,671,042 predicted genes (mean 321,864 genes/fish), of which 1,569,102 were non-redundant at a 90% identity threshold. Estimated gene richness values (Figure 1A) indicated that these samples have likely been sequenced to saturation. Approximately ∼67% (1,044,841) of non-redundant IGC proteins could be assigned a functional homolog in the KEGG, EGGNOG, or Pfam protein family databases (Figure 1B). Similarly, 28% (439,547) of all predicted non-redundant zebrafish microbiome proteins had no homolog in the RefSeq bacteria and archaea protein databases indicating substantial novel protein diversity is encoded in the zebrafish gut microbiome. Of the genes that were annotated, many are predicted to be involved in essential metabolic functions (e.g., sugar, fatty acid, and amino acid metabolism), biosynthesis of short-chain fatty acids, and environmental sensing (Figure 1C). The zebrafish gut microbiome also encoded over 50,000 genes putatively involved in the production of secondary metabolites such as antibiotics, terpenes, and toxins, indicating that the zebrafish gut microbiome may represent an important source of natural product discovery.

**Figure 1.**
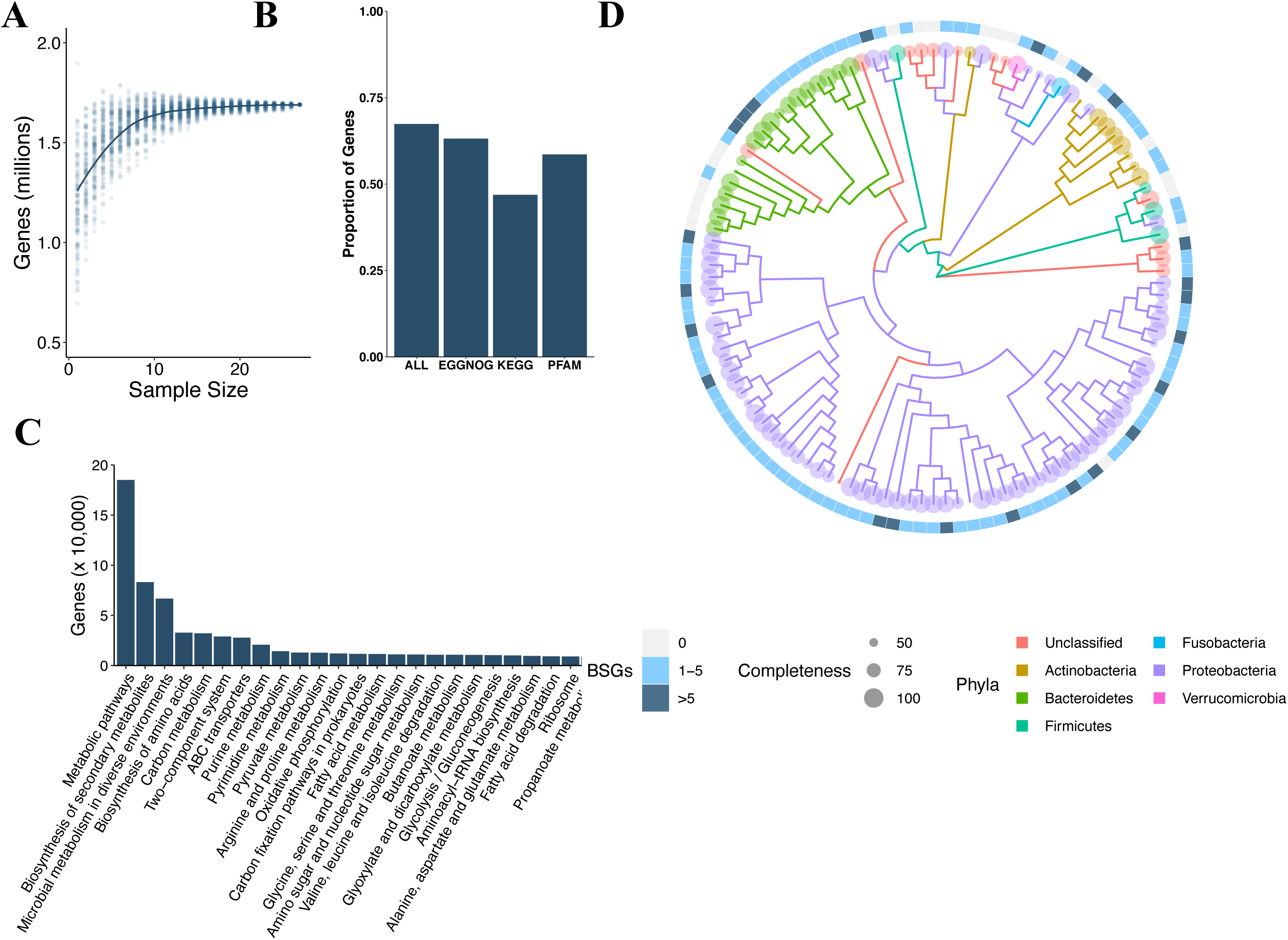
The zebrafish gut contains substantial protein diversity. **A)** rarefaction curve of zebrafish IGC gene richness. **B)** Proportion of zebrafish IGC proteins with a homolog in a any protein family database examined. **C)** The top twenty-five metabolic pathways in the zebrafish integrated gene catalogues. **D)** Phylogenetic tree of genome bins, with terminal nodes labeled.

The products encoded by biosynthetic gene clusters (BGCs), groups of genes that collectively encode a single secondary metabolite biosynthetic pathway, mediate many of the interactions between the microbiome and its environment/host^31^. Systematic investigations of integrated gene catalogs and their underlying metagenomic assemblies have facilitated the identification of substantial novel biosynthetic diversity in the human microbiome^39^ suggesting that host associated microbiomes may encode previously unappreciated BGC diversity. Given the limited knowledge of zebrafish metagenomic diversity, we investigated the identity, novelty, and phylogenetic distribution of BGCs in the zebrafish gut microbiome by sorting assembled metagenomic contigs into 148 genomic bins using an automated expectation-maximization algorithm^27^. Approximately one-third of these bins (48) were estimated to be high quality nearly complete genomes based on marker gene analysis (> 90% complete and < 10% redundant single copy number marker genes; Figure 1D). Across all bins, we found a total of 440 potential BGCs (Supplemental Figure 2A). Less than 50% of proteins present in these clusters share high levels of sequence identity (>80% identity) with RefSeq proteins and ∼15% have no RefSeq homologues (Supplemental Figure 2B). These results collectively indicate that the zebrafish gut microbiome encodes a large diversity of BGCs and many of the genes encoded in these BGCs are likely novel. While the function of the metabolites encoded in these BGCs remains unclear, the observation that microbial natural products interact extensively with microbial and host physiology^39–41^ raises the possibility that some of these compounds may represent drug leads that could benefit human health.

**Figure 2.**
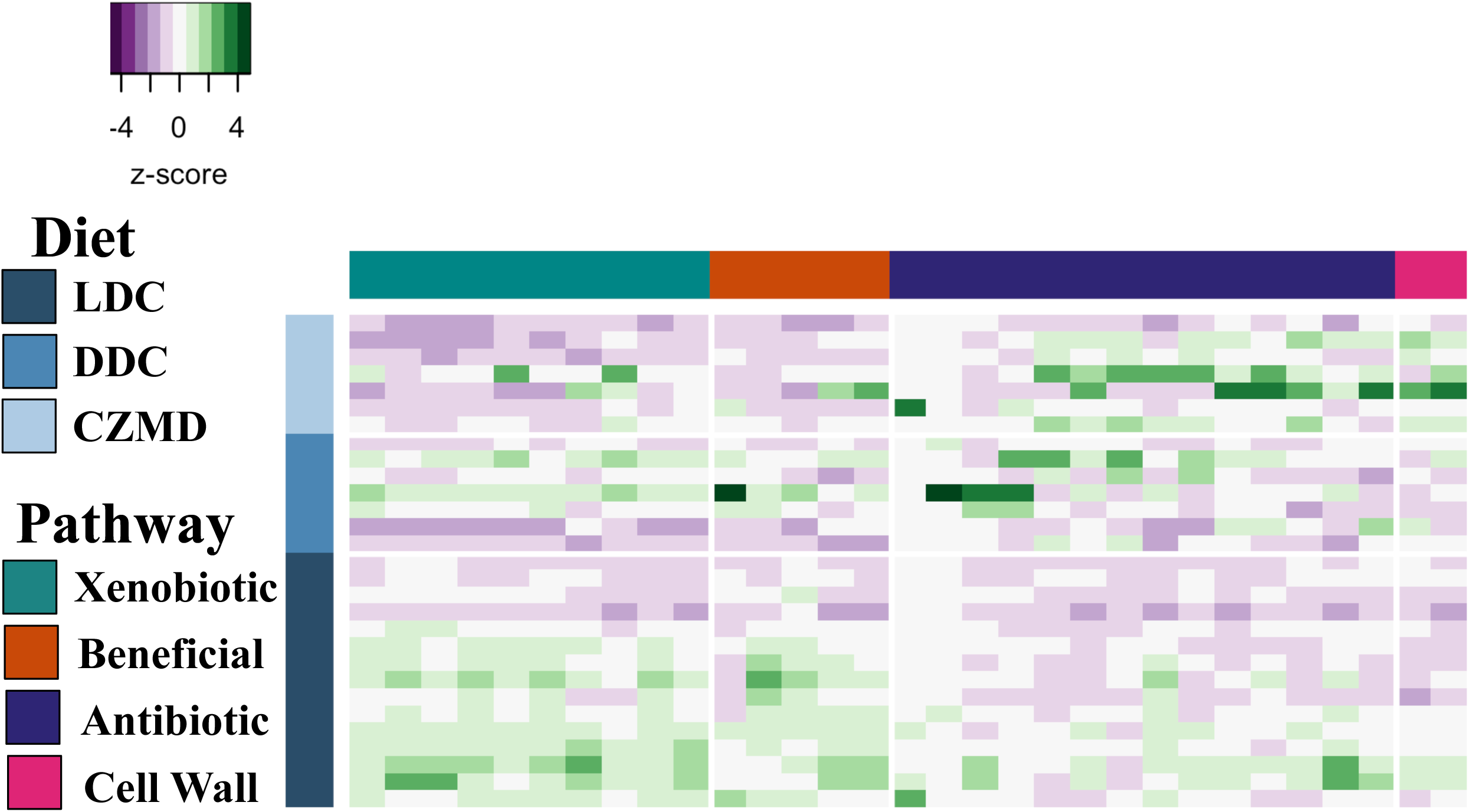
Zinc restriction associates with altered microbiome functional potential. A heat map of select KEGG functional pathways (columns) that vary significantly (q < 0.2) across diets (rows). Row side colors indicate diet. Column colors indicate broad functional groupings of KEGG pathways xenobiotic metabolism (green), beneficial functions (orange), antibiotic production (purple), cell wall (pink).

Since most investigation of microbiome diversity in zebrafish has been limited to 16S or culture-based approaches, we asked if our metagenomic datasets could identify novel microbial diversity that may have been missed by previous surveys. Similar to 16S rRNA surveys ^8^, metagenomic profiling of the zebrafish gut microbiome identifies Proteobacteria, Actinobacteria, Firmicutes, Fusobacteria, and Verrucomicrobia as the dominant phyla in these communities (Supplemental Figure 3A). At finer levels of taxonomic resolution, a substantial proportion of the bacterial species detected (∼41%) were present in all individuals (Supplemental Table 2) supporting previous observations of a robust core microbiome in zebrafish^8^. Taxonomic profiling also detected viral signatures (Supplemental Figure 3B), several of which were ubiquitously present in all samples (Supplemental Figure 3C). Two DNA viruses, Cyprinid herpesvirus 1 and Ictalurid herpesvirus 1, known to infect teleost fish species closely related to zebrafish were amongst these ubiquitous signatures. This observation was consistent with our finding that ∼1% (15,192) of all predicted proteins in the IGC had homologues in eukaryote viruses including several in the family Herpesviridae. Zebrafish have been used as models of herpes virus pathogenesis indicating their susceptibility^42^, however, only one zebrafish virus^43^ and no herpes viruses are known to naturally infect zebrafish. Our findings indicate that viral diversity in the zebrafish may be greater than previously appreciated. If these infections are confirmed, the zebrafish may prove a valuable model for the study of host-virome interactions.

**Figure 3.**
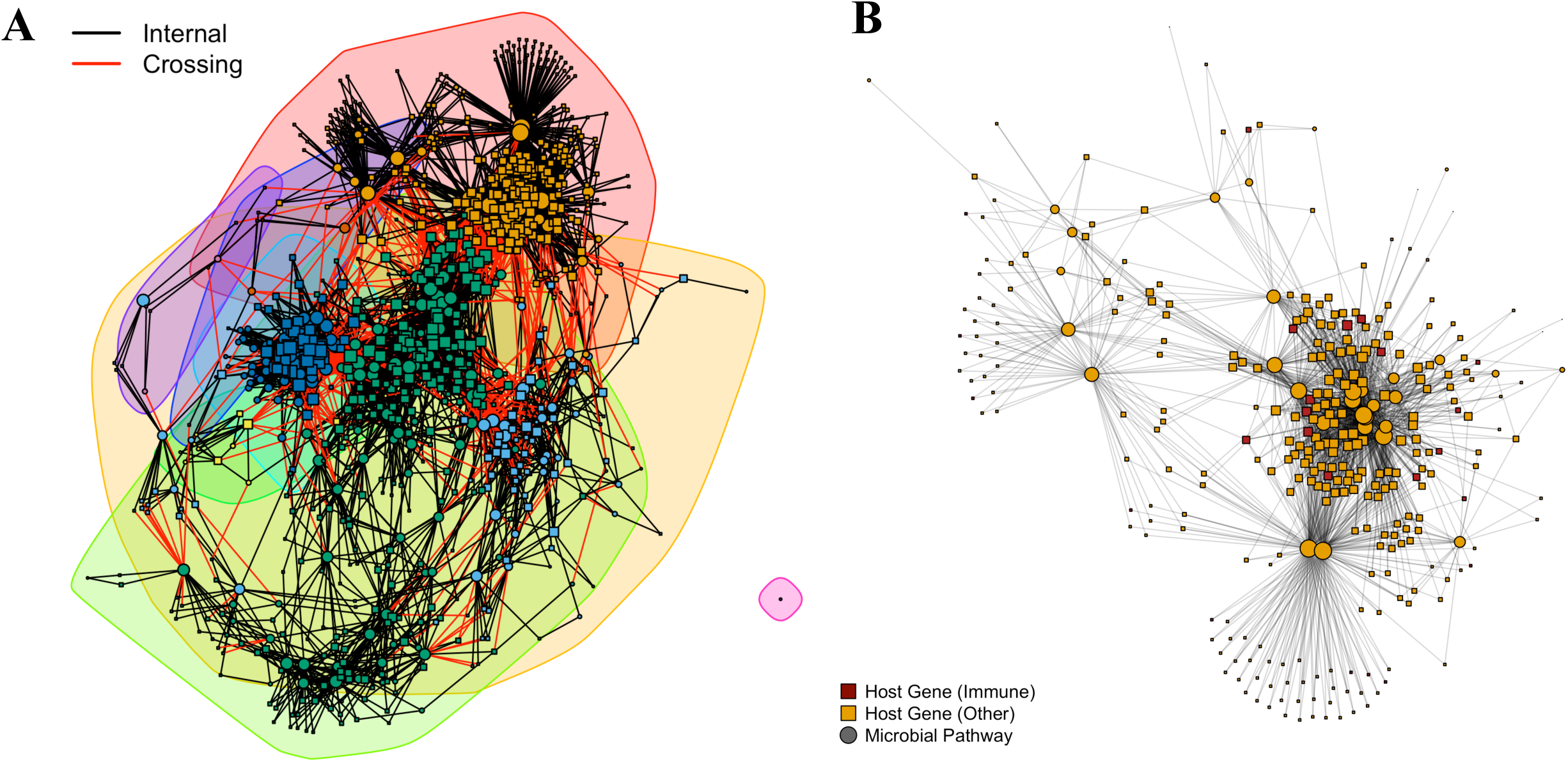
Alteration in gene expression in the gut during zinc restriction correlates with microbiome functional potential. **A)** a correlation network between microbiome genes (circles) and human RNA (squares). Colored ellipses denote network communities. Lines that connect the nodes of the graph represent significant correlations between host RNA and microbiome gene within (black) and between (red) communities. B) A sub graph of vertices that contain highly connected network nodes.

Diet drives microbiome diversity in fish and other vertebrates ^44,45^ and varies widely amongst zebrafish facilities^46^. We used the zebrafish IGC to quantify differences in microbiome functional potential in two lab diets commonly used at the Sinnhuber Aquatic Research Laboratory: a lab diet (LDC) and a purified defined diet (DDC), which differed in macro and micronutrient concentrations (Supplemental Table 1). We also investigated A third diet (CZMD) that was identical to the DDC diet except that the concentration of the essential micronutrient zinc was reduced to levels that induce marginal zinc deficiency in zebrafish^13^. Specifically, we asked how the functional potential of the microbiome varied by macronutrient (i.e., DDC vs LDC) as well as micronutrient concentrations (DDC vs CZMD). Diet did not significantly affect the number of reads that mapped to the IGC (*P* = 0.08), gene richness (*P* = 0.39) or entropy (*P* = 0.28), but did significantly associate with the composition of genes encoded in the metagenome (F_2,26_ = 10.154, R^2^ = 0.44, *P* = 2e-04; Supplemental Figure 4A-E). The abundance of 284 KEGG pathways varied significantly across diets (q < 0.05, Supplemental Table 3). Specifically, the abundance of pathways involved in amino acid metabolism, fatty acid metabolism, and bacterial motility and chemotaxis were elevated in animals fed LDC diet when compared to DDC fed animals, while pathways involved in carbohydrate metabolism and antimicrobial biosynthesis were decreased (Figure 2). These patterns are consistent with high concentrations of protein and low concentrations of carbohydrates in the LDC diet compared to the defined diets. The microbiomes of animals fed zinc restricted diets exhibited lower abundance of pathways associated with beneficial functions including detoxification of xenobiotic compounds and bile acid metabolism when compared with the microbiomes of animals fed the DDC diet. Animals fed the CZMD diet also had elevated abundance of pathways involved in the production of antimicrobial compounds and the biosynthesis of peptidoglycan and lipopolysaccharide (Figure 2). Together these data suggest that dietary variation in macro- and micronutrient composition may modulate how the zebrafish gut microbiota interacts with one another as well as host physiology.

**Figure 4.**
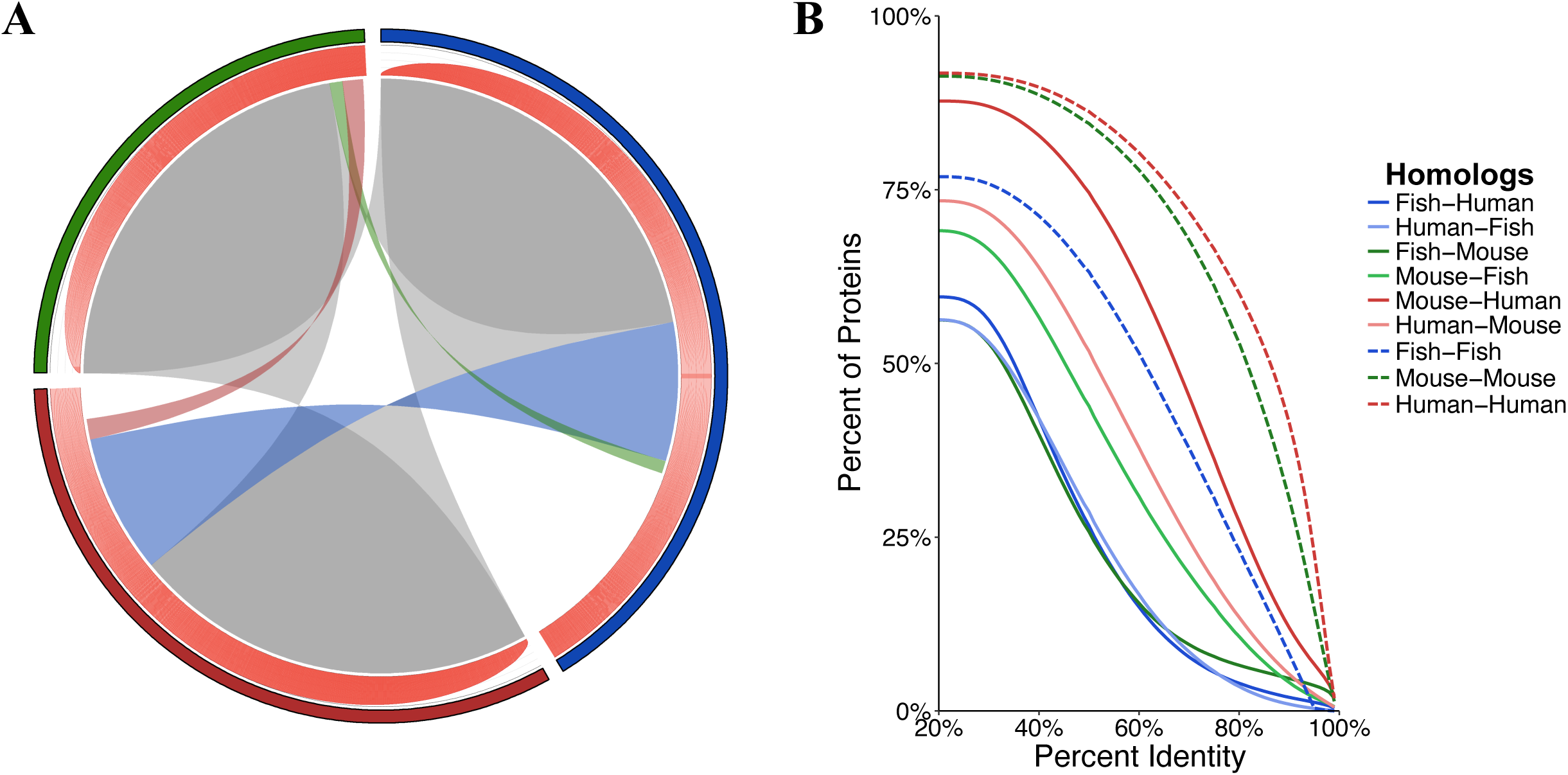
Zebrafish and mammalian microbiomes share significant homology. **A)** A circos plot of KEGG orthologous protein families (outer circle) in zebrafish (blue), humans (red), and mice (green) and their conservation (links) and abundance (inner histogram). **B)** Percentage identity of microbiome proteins between (solid) and within species (dashed). Lines are colored by species pairs.

Dietary variation in zebrafish significantly impacts fecundity, growth, and intestinal inflammation^47–49^, however, the mechanisms underpinning these effects are incompletely resolved. To determine if the observed functional differences in the microbiome across diets associate with altered host physiology, we quantified the effect of diet on whole gut gene expression. Zebrafish intestinal gene expression profiles were significantly stratified by diet (e.g., lab vs defined; F_(2,18)_ = 1.47, R^2^ = 0.14, *P* = 0.004) but not by dietary zinc concentration (F_(1,19)_ = 1.28, R^2^ = 0.06, *P* = 0.080) (Supplemental Figure 5A). In total of 1,315 genes were differentially expressed across diets (q < 0.2, Supplemental Figure 5B). The expression of genes involved in cellular homeostasis, wound healing, and drug metabolism, was lower in animals fed the CZMD diet while animals fed the lab diet had lower expression of genes involved in zinc ion transport and high expression of genes involved in translation and drug metabolism compared to DDC animals.

To determine if variation in microbiome functional potential associates with these changes in host gene expression we quantified correlations between intestinal gene expression and microbiome functional pathway abundances. The resulting 5,978 correlations between 853 zebrafish genes, and 244 microbiome pathways were used to construct a bipartite network (Figure 3). Microbial pathways associated with xenobiotic metabolism, cell wall biosynthesis, and cellular defense response linked extensively with host physiology. Interestingly, these links included associations between microbial pathways and host genes with functions known to be altered during zinc restriction, such as inflammation^50^, wound healing^51^, and DNA damage repair^52^, indicating that the microbiome may be affected by or contribute to these alterations in host physiology. To gain a better understanding of which microbial pathways associate with host functions, we used network community detection analysis, which identifies groups of densely connected network features that may be involved in similar biological processes^53,54^. One of the largest sub-networks was comprised of 316 nodes and included highly connected microbiome pathways associated with bacterial stress response and defense streptomycin biosynthesis pathway, which are increased in zinc-restricted animals. For example, the streptomycin biosynthesis pathway (ko00983) linked to host genes that are involved in immune response, apoptosis, and DNA damage repair. Similarly, the drug metabolism (ko00983) pathway, which is involved in biodegradation and metabolism of xenobiotic compounds, including antibiotics was also a highly connected member of this sub-network that linked to similar host genes. While it remains unclear if the changes in pathway abundance are a cause or consequence of altered host gene expression, these results raise the possibility that the microbiome may contribute to host physiological changes associated with zinc restriction.

As human microbiome research transitions from cataloguing microbial diversity to understanding the mechanisms that underpin host-microbiome interactions, animal models that consistently yield translationally relevant results are crucial. We assessed the translational potential of zebrafish gut microbiome research by quantifying similarities between zebrafish and human gut microbiomes. Similarities were also assessed between zebrafish and mouse gut microbiomes as mice represent the most frequently used, and putatively translationally relevant, model of human microbiome operation. Consistent with previous reports, human and mouse gut microbiomes are composed largely of Firmicutes, Bacteroidetes, and Verrucomicrobia while the zebrafish gut microbiome is composed of Proteobacteria and Fusobacteria (Supplemental figure 6A). However, the relative proportions of zebrafish IGC genes involved in specific metagenome functions (i.e., KEGG pathways) was similar to the proportions in humans (ρ = 0.92 P < 2.2×10^− 16^) and mice (ρ = 0.91 P < 2.2×10^−16^) suggesting that there may exist broader functional overlap between fish microbiomes and that of other vertebrates than the incongruity of their taxonomic composition suggests.

We reasoned that measuring this overlap would provide insights into the potential translatability of zebrafish microbiome research. Despite consisting of fewer individuals and genes, the zebrafish IGC was surprisingly diverse when compared to that of the IGCs of mice and humans. For example, zebrafish had higher KO richness and Shannon entropy (Supplemental Figure 6B-D), more unique functional annotations to individual KEGG, EGGNOG, and Pfam protein families (Supplemental Figure 6E), and a higher percentage of database protein family coverage than did mice and humans (Supplemental Figure 6F). Despite greater functional diversity in the zebrafish IGC, fish, humans, and mice share > 50% of their KO diversity and the vast majority (> 99%) of an individual sample’s KO abundance is contained in this shared fraction (Figure 4A). This observation further suggests that zebrafish and mammalian microbiomes may execute similar functions. A smaller fraction of KO diversity is shared exclusively between two host IGCs or uniquely encoded in one host IGC, indicating that some potential host-microbe functional interactions may be host lineage specific.

Since only a fraction (47-63%) of predicted microbiome protein diversity can be functionally annotated in databases, and since database based classification methods are limited by database size, scope, and maintenance^55^, we next measured homology among predicted IGC proteins across host species. Over half of the predicted zebrafish microbiome proteins had a mouse (56%) or human (59%) microbiome homolog (Figure 4B). Mammalian microbiomes shared more homologues (73%-88%) with one another than they did with fish microbiomes (56%-69%). Human and mice IGCs also had higher levels of intra-IGC homologs (91% and 92%, respectively) than zebrafish (77%), which could reflect that a more diverse collection of proteins are present in the zebrafish IGC or that human and mouse metagenomic diversity has been more exhaustively sequenced.

The taxonomic disparities between humans and zebrafish gut microbiomes make it unlikely that microbes that associate with a physiological phenomenon in fish will be present at appreciable levels in the human gut microbiome. However, given the extensive homology between predicted proteins from human, mouse, and fish gut microbiomes, it is possible that microbes may be present in the mammalian gut microbiome that encode genes that elicit similar effects. We reasoned that by grouping human, mouse, and zebrafish proteins into protein families we could create a database of protein homologs that could enable rapid identification of translationally relevant discovery. For example, if a zebrafish gut microbiome protein associates with intestinal inflammation we could identify all human, mouse, and zebrafish proteins that are closely related to this protein by identifying the family to which the protein of interest belongs. We could then use this information to design experiments that would explicitly test if these homologous proteins associate with similar effects in mice and humans. We generated a network of 2,815,252 protein families comprised of 13,742,715 predicted proteins from zebrafish (1,451,303), mouse (2,544,637), and human (9,746,775) IGCs (Supplemental Figure 7A). While only a small fraction of these protein families (32,470) were universally shared (*i*.*e*., contained proteins from zebrafish, mouse, and human IGCs), the majority of individual proteins (53%) in this network were contained within these universally shared families. The majority of protein abundance in zebrafish (53%), mice (93%), and humans (81%) also was contained within the universally shared families together indicating the tremendous diversity and potential importance of these families. Interestingly, over half (19,634) of the universally shared protein families exhibited patterns of phylogenetic association with host species evolutionary distance. While the mechanisms that underpin these patterns are unclear at the level of microbiome function, co-diversification, environmental filtering, and co-evolution^56^ are hypothesized to drive similar associations between bacterial taxonomy and host lineage.

To evaluate the ability of this vertebrate microbiome protein network to distinguish translationally relevant discoveries in fish, we identified protein families that contained homologs to Beta cell expansion factor A (BefA), a zebrafish microbiome protein associated with pancreatic Beta cell development^57^. Consistent with the findings of Hill and colleagues, we successfully identified ten potential distant human BefA homologs (20-25% identity) in two clusters using this approach. Interestingly, these two clusters contained only zebrafish and human proteins but no mouse homologs, suggesting that there are some functional interactions between vertebrates and their gut microbiota that may not be represented in the mouse model. To confirm this result, we then aligned BefA to all proteins in the mouse IGC and, despite using permissive thresholds (>20% a.a. identity, e < 0.001), were unable to identify BefA homologs in mice. While interactions between BefA and host physiology have been established in zebrafish, the role of this protein in microbiome-host interactions in mammals is unclear. It is possible that BefA homologs do not associate similarly with mammalian physiology and thus have been lost in mice. Regardless, these observations encourage the use of caution when selecting animal models for microbiome-based experiments and necessitate analyses with a broader phylogenetic scope of hosts aimed at evaluating the translational relevance of animal models on a protein-by-protein basis.

As the use of zebrafish in microbiome research increases, the IGC generated here will enable the identification of the genomic underpinnings of host-microbiome interactions in zebrafish. While the finding that most functional diversity is shared between humans, mice, and zebrafish supports the use of this system for study of microbiome operation, the modest levels of identity between zebrafish and human homologs dictates that care be used in the evaluation of the translational relevance of findings in this model. The vertebrate protein network produced here will provide a resource for researchers to rapidly perform those translational assessments. Moving forward, the incorporation of metagenomic data from additional zebrafish facilities, strains, and environmental conditions will continue to expand our understanding of zebrafish functional diversity and will likely reveal other areas where the zebrafish may be well suited to investigate specific host-microbiome interactions.

## Acknowledgements

The authors thank the member of the Center for Genome Research and Biocomputing at Oregon State University for their assistance with sequencing and maintenance of our computational infrastructure and the staff of the Sinnhuber Aquatic Research Laboratory for assistance with animal care and husbandry. This work was supported by awards from the National Institutes of Health (P30-000210, R21-ES023937, 5R21-AI135641), the United States Department of Agriculture (NIFA/USDA-2018-67017-27358) and the National Science Foundation (award 1557192).

---

**Supplemental Figure 1.**
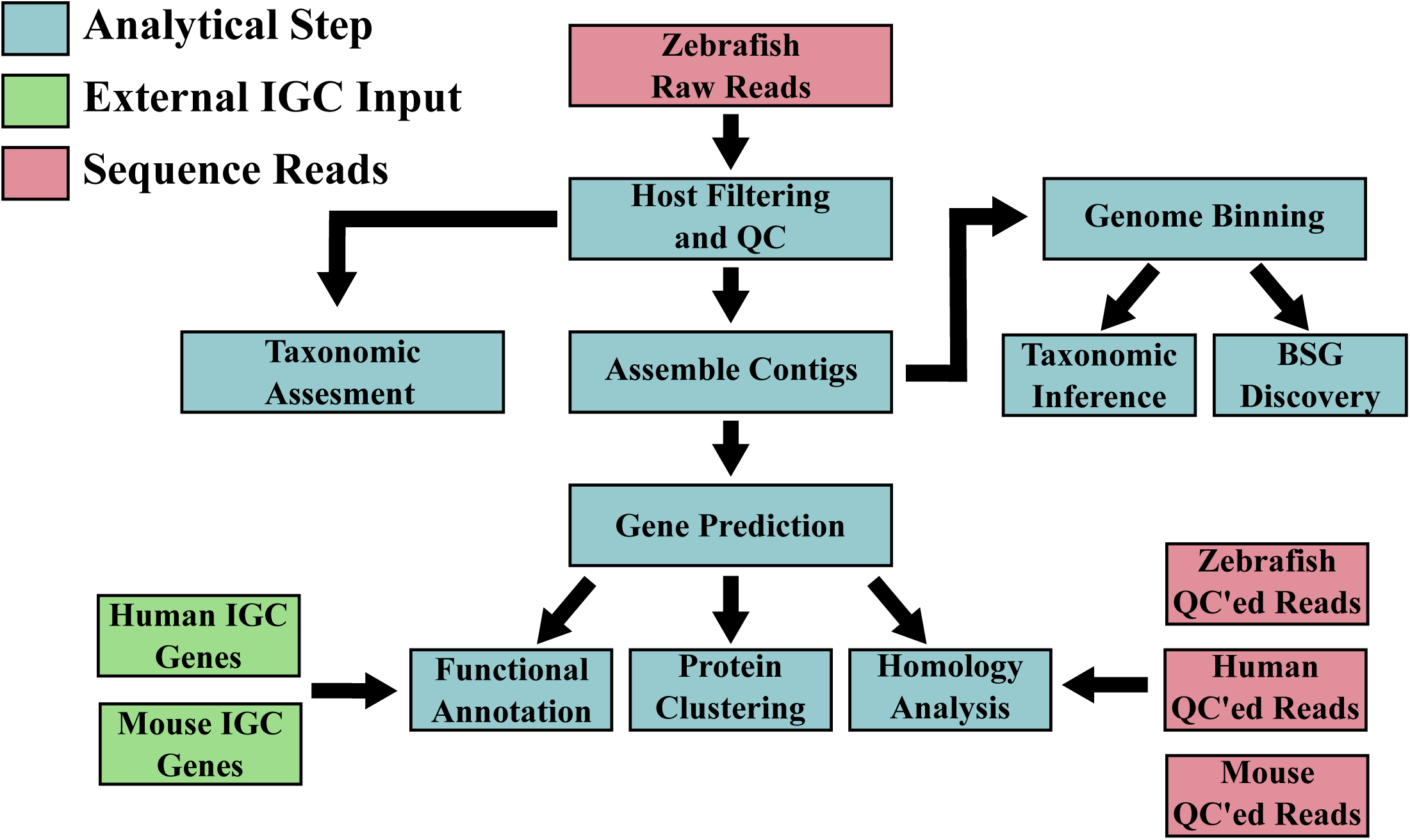
Workflow diagram for zebrafish integrated gene catalogue construction and downstream analysis. Each analytical step of the analysis (blue) and areas where genes from other integrated gene catalogues (IGC; green) or sequence reads were incorporated (red) are illustrated.

**Supplemental Figure 2.**
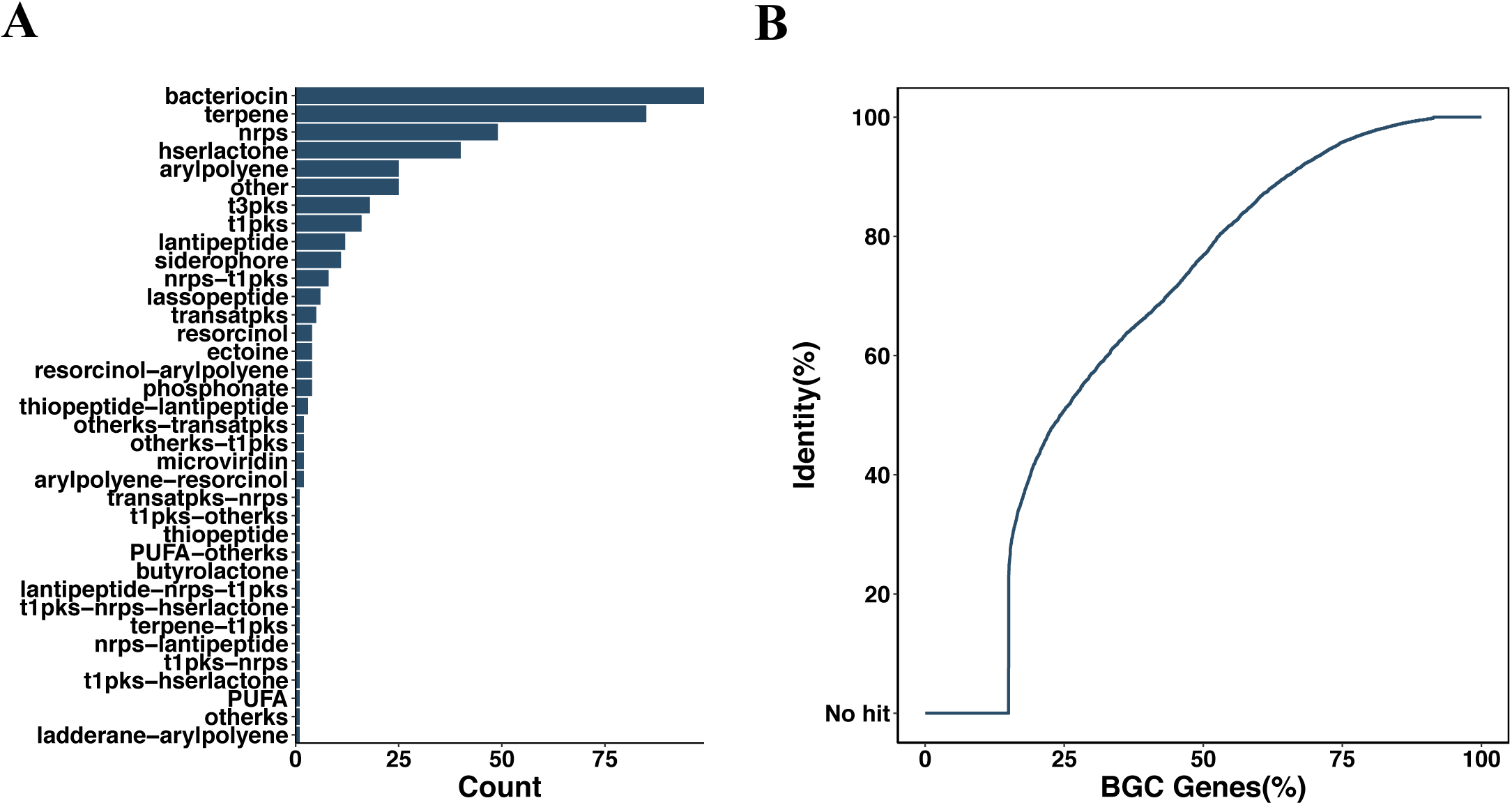
Zebrafish encode diverse natural product diversity. **A)** bar chart of predicted function of biosynthetic gene clusters found in the zebrafish gut microbiome. **B)** Line graph depicting the number of biosynthetic gene cluster protein with RefSeq homologs at a give identity threshold.

**Supplemental Figure 3.**
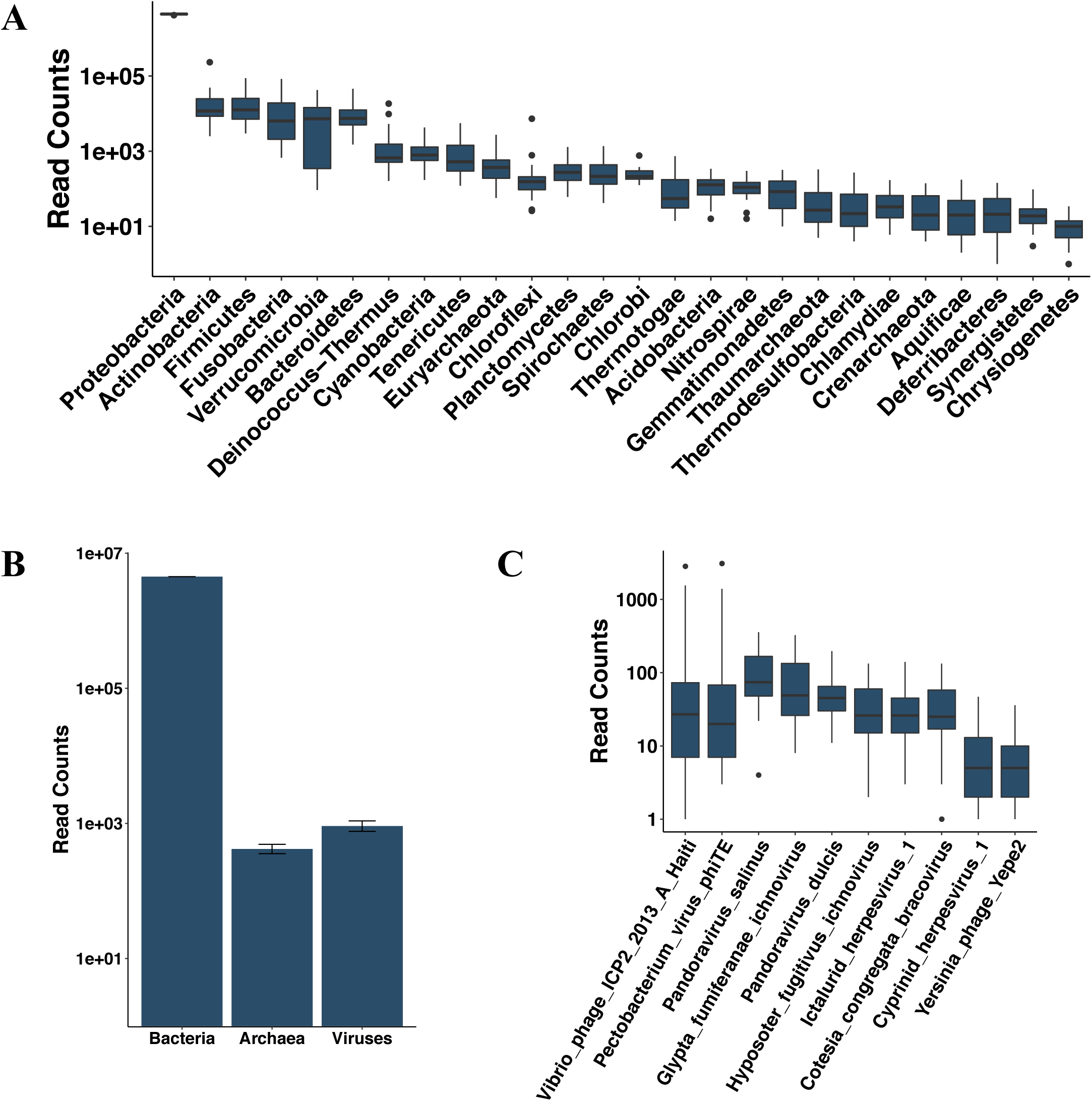
Novel insights into the zebrafish gut microbiome’s compositional diversity. **A)** Abundance of bacterial phyla in the zebrafish gut microbiome. **B)** Number of reads from the zebrafish gut microbiome that associate with bacterial, archeal, and viral taxa. **C)** Zebrafish gut microbiome read counts associated with ubiquitous viral taxa.

**Supplemental Figure 4.**
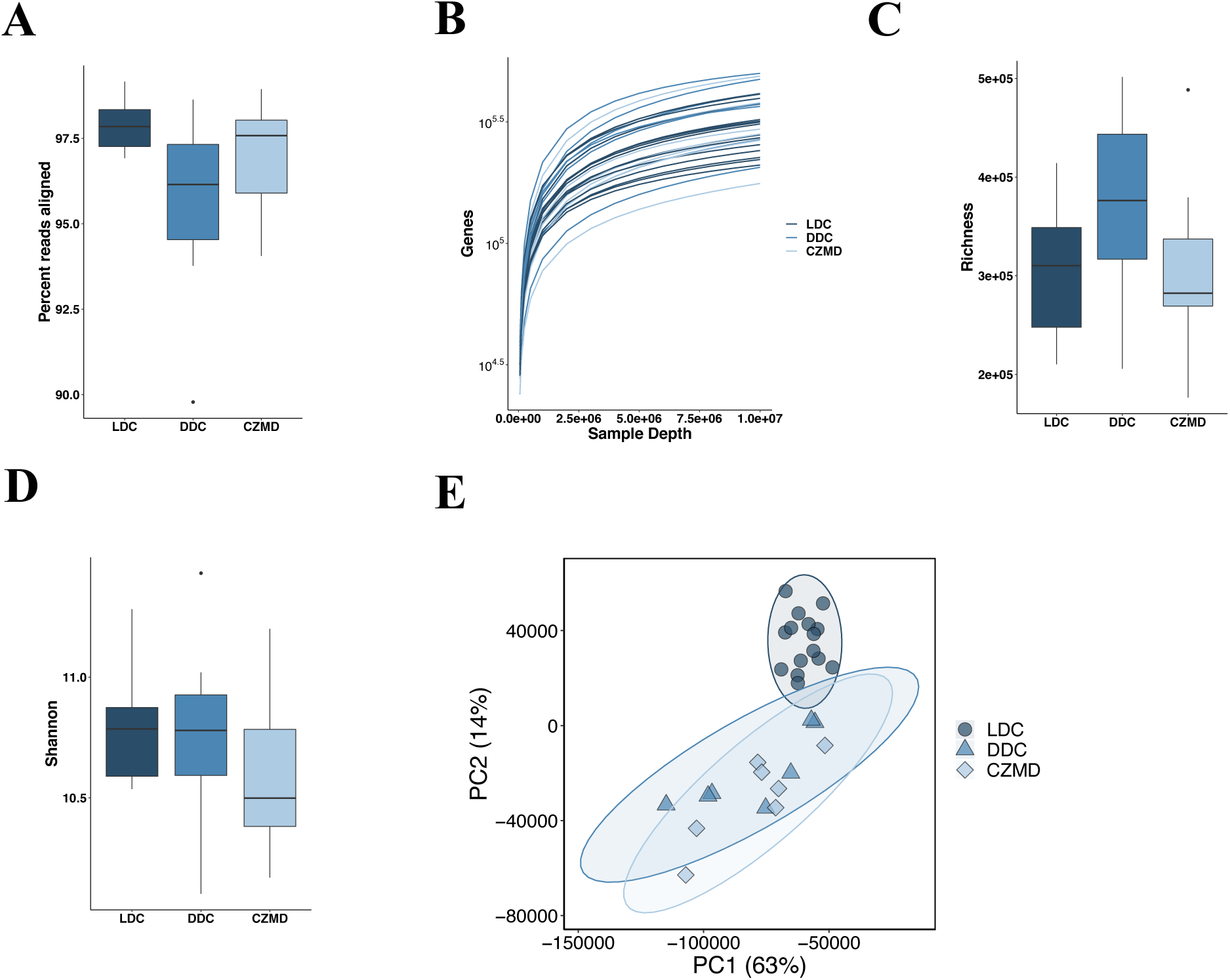
Diet significantly impacts zebrafish microbiome function. **A)** percent read that mapped to zebrafish IGC protein models by species. **B)** Gene rarefaction by diet. **C)** boxplots of gene richness and **D)** Shannon entropy colored by diet. **E)** Principal component plot of zebrafish microbiome genes.

**Supplemental Figure 5.**
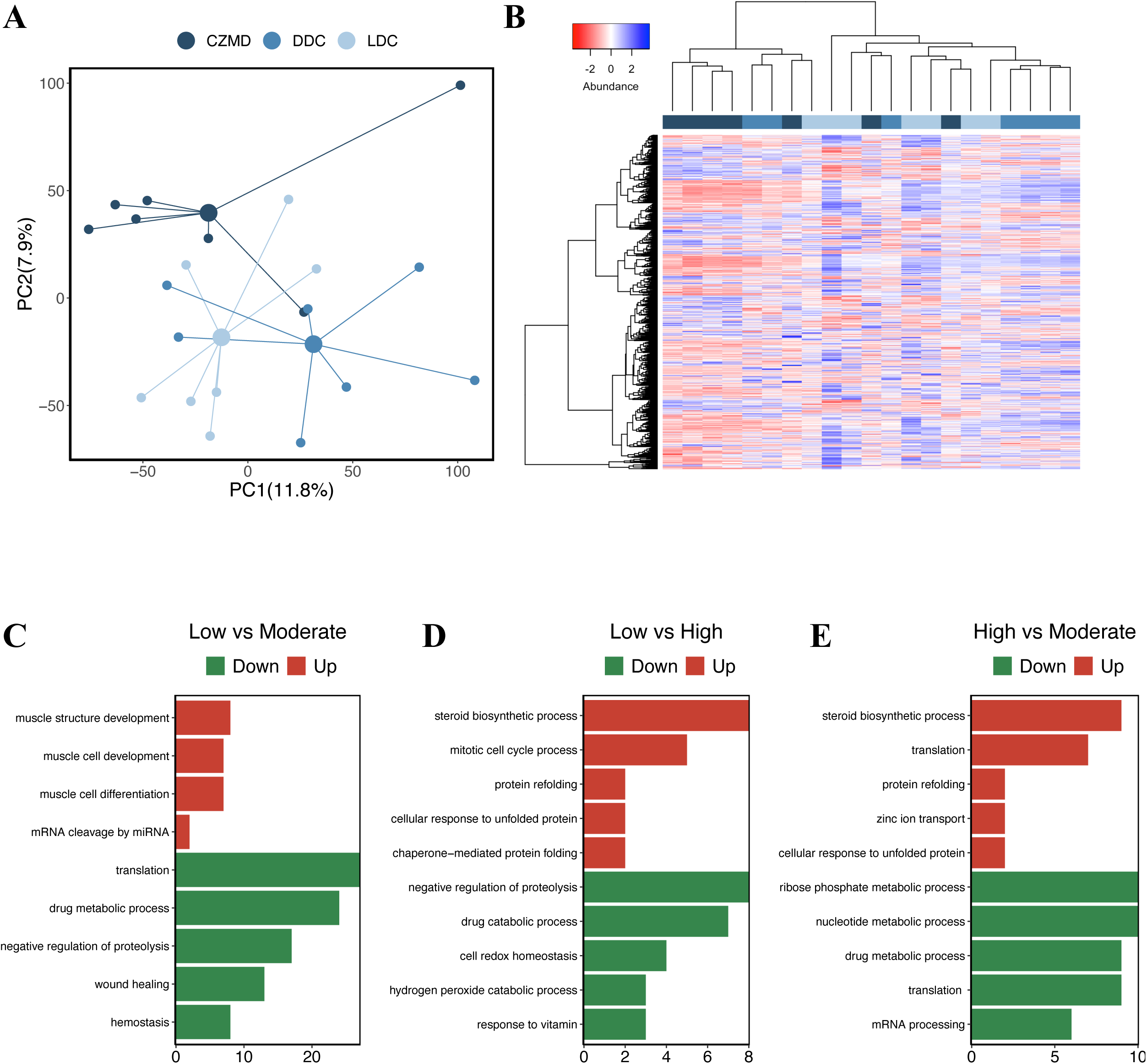
Zinc restriction alter zebrafish gut gene expression. **A)** Principal components plot of zebrafish gut gene expression by diet. B) Heat map of gene expression scaled by row. Column colors of the heat map indicate CZMD (red), DDC (green), or LDC (blue) diets. **C-E)** Selected GO terms that are enriched (q < 0.2) in various comparisons of diet. Color indicates the direction of gene expression change.

**Supplemental Figure 6.**
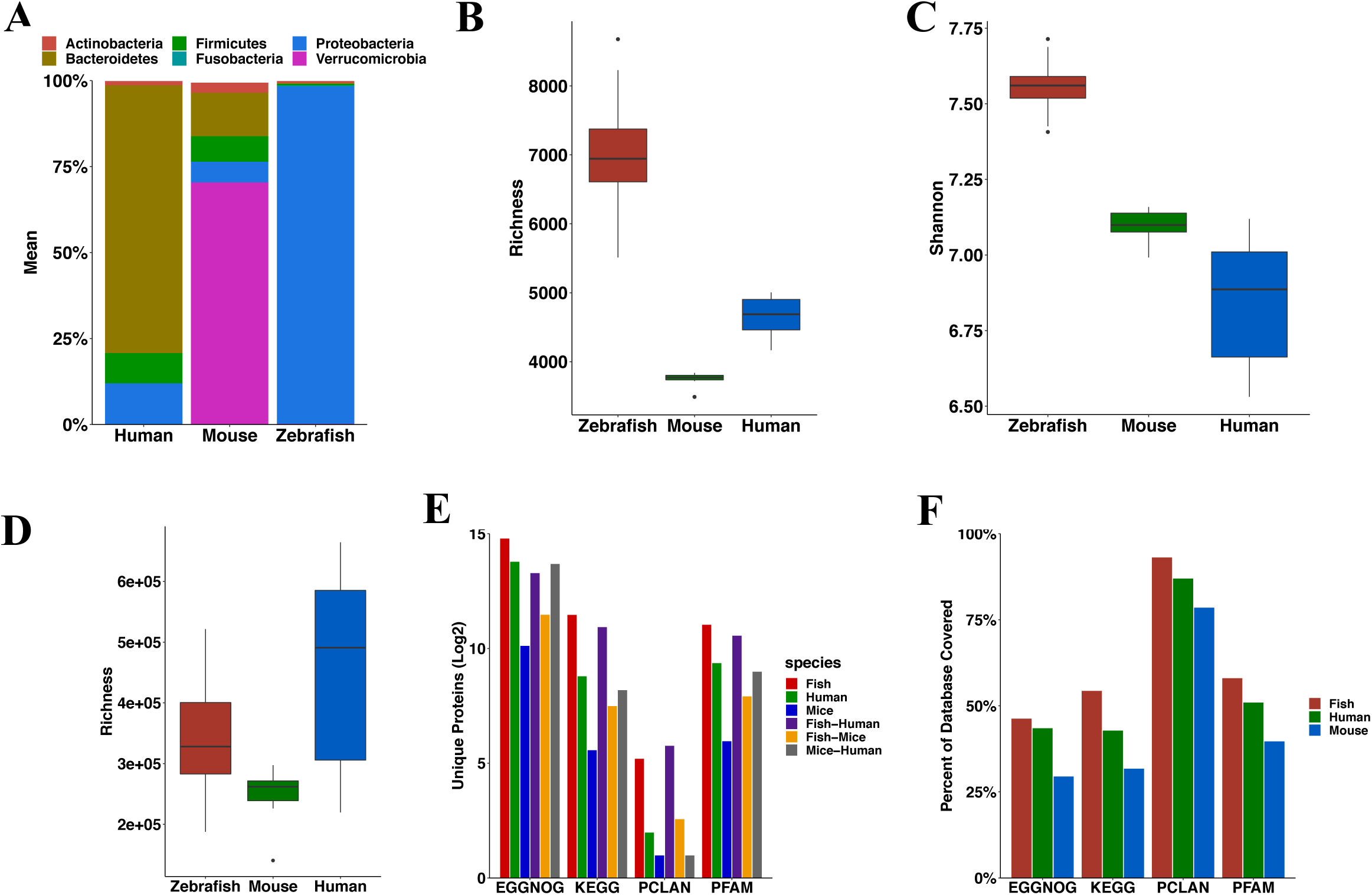
Zebrafish microbiomes more diverse that mice and humans. **A)** The ten most highly abundant phyla in human, mouse, and fish microbiomes **B)** KEGG KO richness **C)** Shannon entropy boxplots by species. **D)** Gene richness by species. **E)** The number of hits to protein databases by species. **F)** Percent protein database coverage by across species.

**Supplemental Figure 7.**
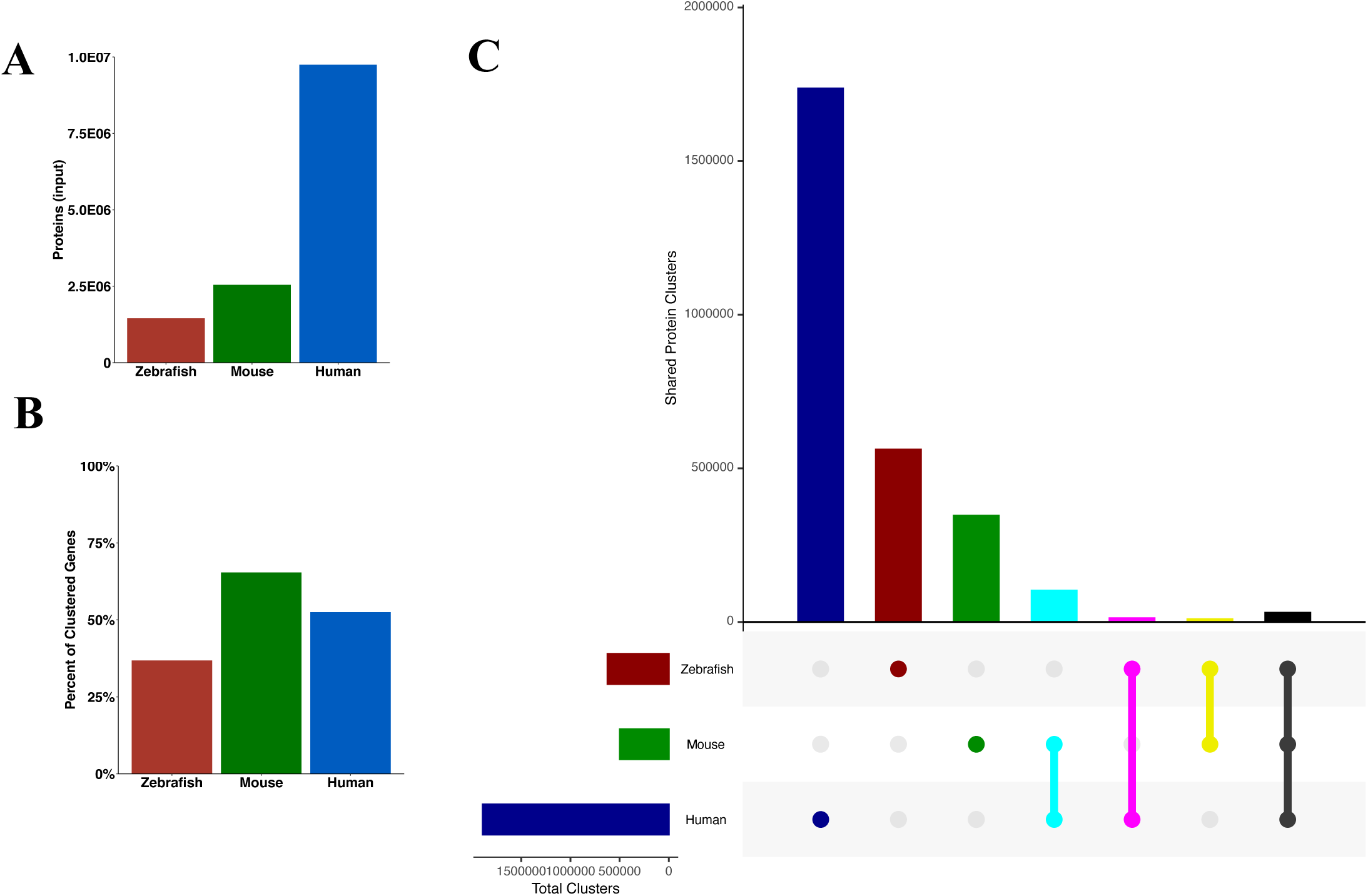
Substantial protein family diversity is shared between humans, mice and zebrafish. **A)** Total number of proteins clustered in MCL analysis. **B)** Percent of proteins clustered that are present in clusters that are universally shared across species. **C)** Bar plot depicting unique and shared protein diversity in each species.

